# Anti-nuclear antibody (ANA) detection in culture supernatant of cloned B cells

**DOI:** 10.1101/2023.05.12.540540

**Authors:** Shabirul Haque, Manami Watanabe, Yemil Atisha-Fregoso, Betty Diamond

## Abstract

Anti-nuclear antibodies (ANA) of individuals with SLE contribute to disease pathogenesis and tissue injury. Systematic analysis of the fine specificity of ANA produced by individual B cells has been limited, in part, to the low concentration of immunoglobulin in single B cell cultures. We modified single B cell culture methods and developed an ANA ELISA to detect ANA at very low levels (15.0 ng/ml) of IgM in the supernatant of clonally expanded B cell cultures. This approach allows for the analysis of ANA reactivity of the IgM produced by approximately 60% of the single sorted naive B cells expanded in culture and permits a relatively high throughput evaluation at the single cell level of the ANA reactivity of naïve B cells of healthy individuals and patients with autoimmune diseases.

## 1. Introduction

In normal individuals more than 50% of the early immature B cells that are generated by random rearrangements of VDJ segments show some degree of autoreactivity [1]. These cells are partially removed by discrete immune checkpoints, however, it has been shown by us [2] and others [1] that approximately 20% of the mature naïve B cell compartment is comprised of autoreactive B cells.

High titers of autoantibodies against nuclear antigens (ANA) are present in a variety of autoimmune conditions such as systemic lupus erythematosus (SLE) [3-5], scleroderma [6], polymyositis, autoimmune hepatitis [7], drug induced lupus [8] and mixed connective tissue disease [9]. In some cases, they are considered markers of disease, but in others they are considered pathogenic. The defects that cause increased titers of these antibodies have been extensively studied. There is evidence that the underlying alteration that leads to the production of pathogenic antibodies in patients with SLE is a defect in the negative selection of autoreactive B cells at the time of establishing the repertoire of immunocompetent B cells. Naïve autoreactive B cells are then able to be activated and participate in autoantibody production [10,11]. There is also evidence for normal frequency of autoreactive B cells in the naïve B cell compartment but increased B cell differentiation to plasma cells. [12].

An analysis of the ANA produced by naïve B cells addresses issues regarding the selection of autoreactive B cells. We previously developed a method for the identification of ANA+ B cells by flow cytometry. Here we describe a method to expand single sorted naïve B cells and to detect ANA in culture supernatants. This will permit a comparison of ANA+ B cells in healthy individuals and those with an autoimmune disease and comparison of ANA+ B cells across autoimmune diseases.

## 2. Materials and methods

### 2.1 Reactants and materials

Culture plate (96 well culture plate, ref# 353072, treated by vacuum gas plasma polystyrene non-pyrogenic, Falcon). ELISA plate (costar assay plate, 96 well clear, flat bottom, half area high binding, polystyrene, cat# 3690, Corning). Streptavidin-UNLB (cat # 7105-01, Southern Biotech). Anti-dsDNA monoclonal antibody (G11) and a control monoclonal antibody (B1) was developed as described [13,2]. Nuclei EZ lysis kit (Sigma NUC-101). EZ-Link-Sulfo-NHS-LC-biotin, (Thermo #21355).

#### B cell medium (BCM) composition

RPMI-1640 (Gibco, 11875-093) supplemented with 10% FCS (Thermo Scientific, SH30070.03 Hyclone), 55 μM 2-ME (Gibco, 21985), 1%Pen Strep (Gibco, 15140-122), 10 mM HEPES (Gibco, 15630-080), 1 mM Sodium Pyruvate (Gibco, 11360-070), 1% MEM NEAA (Gibco, 11140-050).

#### Feeder cells (MS40L-low) medium

IMDM (Invitrogen, 12440-053) supplemented with 10% FCS (Hyclone), 1% Pen Strep (100 Units/ml Penicillin, 100 μg/ml Streptomycin), and 55 μM 2-ME (GIBCO).

#### Antibodies

Goat polyclonal anti-human Ig-UNLB (cat# 2010-01, southern biotech). Goat polyclonal anti-human IgM-UNLB (cat# 2020-01, southern biotech). Human IgG (cat# 0150-01, Southern Biotech) + Human IgM-L (cat# 0158L-01, southern biotech) + Human IgA-k (cat# 0155K-01, Southern Biotech). Goat anti-human IgM-AP (cat# 2020-04, Southern Biotech), goat anti-human IgG-AP (cat# 2040-04, Southern Biotech), and goat anti-human IgA-AP (cat # 2050-04, Southern Biotech).

#### Cytokines

Recombinant human IL-2 (Peprotech 200-02, final concentration for culture is 50ng/ml), Recombinant human IL-4 (Peprotech 200-04, final concentration for culture is 10ng/ml), Recombinant human IL-21 (Peprotech 200-21, final concentration for culture is 10ng/ml), Recombinant human BAFF (Peprotech 310-13, final concentration for culture is 10ng/ml).

### 2.2. Processing of human blood samples

Blood samples were collected from healthy adult donors. Peripheral blood mononuclear cells (PBMCs) were isolated by density gradient centrifugation using Cytiva Ficoll-Paque PLUS (45-001-749, Fisher Scientific).

### 2.3. Preparation and biotinylation of nuclear extract (NE)

Nuclear extracts were prepared as previously described [2]. HeLa cells were grown to confluence in a T75 flask, and nuclei were isolated using a Nuclei EZ lysis kit, according to the protocol of the manufacturer (Sigma-Aldrich). The nuclei were washed with phosphate buffered saline (PBS) and pelleted at 500g for 5 minutes. The pellet was resuspended in 3.0 ml of PBS containing a complete Mini Protease Inhibitor tablet (Roche), split into two eppendorf tubes prefilled with 150 μl of 0.5-mm glass beads (Scientific Industries) in each tube, and vortexed vigorously on a Disruptor Genie for 1 hour at 4 °C. The nuclear extract was then spun at 21,000g for 5 minutes and the supernatant recovered and biotinylated using an EZ-Link Sulfo-NHS-LC-Biotin kit (Thermo Scientific). The solution was dialyzed using Slide-A-Lyzer® Dialysis Cassette G2 (cat# 87724, thermos scientific) in PBS for 24 hours at 4 °C, and aliquots were stored at -80 °C.

### 2.4. Cell staining for flow cytometry

ANA staining of B cells was performed as previously described [4,2]. PBMCs were washed with washing buffer (5% sterile FBS in HBSS filtered), spun down and supernatant was removed. Two uL of the biotinylated-nuclear extract were diluted into 300 μl of 1.5% fat free milk (lab scientific, cat # M0841) in HBSS (filtered with sterile cell strainer, Fisher Scientific, cat# 22-363-548). The solution was vortexed for 30 seconds and added into the washed cells and mixed. The cells were incubated on ice for 30 minutes. After incubation, cells were washed twice with 5.0 ml of washing buffer (300g for 5 minutes at 4°C). Conjugated antibodies were prepared in 300μl staining buffer (CD19-PECy7, 1:50; CD27-PE, 1:50; CD38-PEeF610, 1:50; Streptavidin-APC, 1:200;

FVD-eF506, 1:500) and mixed antibodies were added to the cells which were incubated on ice for 30 minutes in dark. Cells were washed twice, and pellets were re-suspended in 600 μl 5% sterile FBS in HBSS. Re-suspended cells were analyzed and ANA+ and ANA-naïve B cells (CD19+, CD27-, CD38 intermediate) were directly sorted into the culture plate for cell culture.

### 2.5. Single B cell cultures

The culture of single B cells was performed as described [14-16] utilizing the MS40L-low feeder cells previously reported [17]. Briefly, culture plate wells were coated with feeder cells (3000 cells/100 μl/well) 24 hours prior to B cell sorting in B cell medium (BCM) and incubated overnight in 5% CO2 at 37 °C incubator. The next day, 1-2 hour prior to B cell sorting, 100 μl/well, 2X cytokines were added into the culture plate. Either single cell or multiple cells (as a positive control) were sorted into the culture plate and the plate was placed in 37 °C and 5% CO2 incubator. Every 4 days, culture medium was replaced by gently removing 100μl of supernatant and adding 100μl of fresh medium with cytokines. Culture supernatant was collected at day 20 - 28 for assessment of Ig and ANA production.

### 2.6. Measurement of Immunoglobulin (Ig) by ELISA

ELISA plates were coated with 25μl/well goat polyclonal anti-human Ig-UNLB (1:100 diluted in 1x PBS) for total Ig ELISA or goat polyclonal anti-human IgM-UNLB (1:500 diluted in 1x PBS) for IgM ELISA, and the plate was incubated at 4°C overnight with cover. The next day, the plate was washed four times with washing buffer (PBS 0.05% Tween-20) and blocked with blocking buffer (1% BSA in PBS) for 1 hour at room temperature (RT) and plates were washed. Culture supernatants (5-fold diluted) from the B cell cultures were added to the wells (25μl/well). Serially diluted immunoglobulins IgG+IgA+IgM (200, 100, 50, 25, 12.5, 6.3, 3.1, 1.6, 0.8, 0.4, 0.2 and 0 ng/ml) were also added to the plate to generate a standard curve. The plate was incubated for 2.0 hours at RT. The ELISA plate was washed four times. Secondary antibodies (anti-IgG + anti-IgM + anti-IgA for total Ig or anti-IgM for IgM ELISA) conjugated with alkaline phosphatase (AP) were diluted 1:1000 in 0.2% BSA in PBS, mixed and 25μl/well were added to the wells. The plate was incubated for 1.0 hour at RT and washed four times. Phosphatase substrate buffer was added (50μl/well) and the plate was incubated until color developed at 37°C in the dark. Absorbance of each well was measured at 405 nm (Victor3, 1420 multilabel counter plate reader, Perkin Elmer).

### 2.7. Measurement of ANA by ELISA

ELISA plates were coated with 25 μl of unlabeled streptavidin (2.0 μg/ml diluted in distilled water) and incubated at 4°C overnight with cover. The next day, the plate was washed four times with washing buffer (PBS 0.05% Tween-20) and blocked with blocking buffer (1 % BSA in PBS) for 1 hour at 37°C. Biotinylated nuclear extract (Bio-NE 30ug/ml diluted in 0.2% BSA) were incubated for 1 hour at 37°C. After the incubation, the ELISA plate was washed four times. Primary antibodies (culture supernatants) were added 1:1 dilution in 0.2% BSA in PBS. After the 2 hours incubation, the ELISA plate was washed four times. Secondary antibodies (IgM-AP) were diluted 1:500 in 0.2% BSA in PBS and added (25 μl/well) into the plate. The ELISA plate was incubated for 1 hour at RT and then washed four times. Phosphatase substrate buffer was added (50μl/well) and the plate was incubated until color developed at 37°C in dark (approximately 90 minutes were optimum time for color development). The plate absorbance was measured at 405 nm.

### 2.8. Statistical analysis

The cut off for ANA positivity was decided based on the mean of ANA-samples + 2SD. The comparison of OD values was performed using independent samples student T-test.

## 3. Results and Discussion

### 3.1. Ig production in single sorted B cell cultures

Single naïve ANA- and ANA+ B cells were identified by flow cytometry (**Fig. 1A**) and directly sorted into 96 wells culture plates. The gating strategy to identify naïve B cells among live CD19+ B cells is shown (**Supplemental Fig. 1**). Sorted ANA-naïve B cells (82 cells), and ANA+ naïve B cells (82 cells) were cultured for 26 days and the IgM concentration in culture supernatant was measured by ELISA. In 81% of the wells of ANA-naïve B cells and 68% of the ANA+ naïve B cells, we detected production of IgM within the linear range of the assay (above 1.5 ng/mL) (**Fig. 1B**). As shown below, the IgM-ANA ELISA test developed for this assay can detect as little as 15.0 ng/ml in B cell culture supernatant. In total, 57% of the sorted naive B cells produced IgM above the limit of detection of the ANA ELISA.

**Figure 1.**
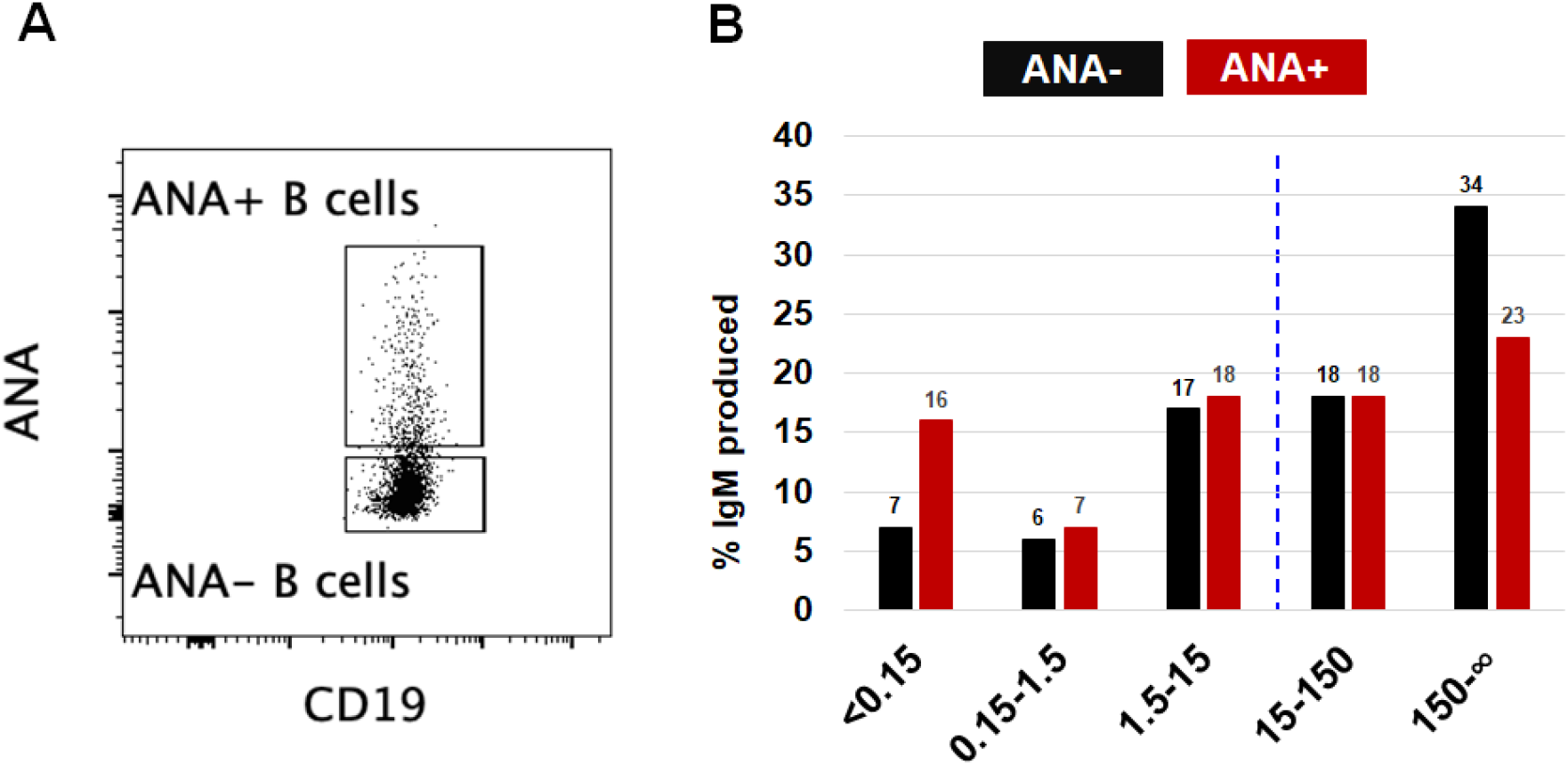
ANA- and ANA+ naïve B cell sort and IgM production. **A**. Gating strategy for ANA- and ANA+ naive B cells. Cells were sorted into 96 well culture plates and kept in culture for 26 days according to the described protocol. **B**. Production of IgM was assessed by ELISA. An IgM concentration >15 ng/ml is required for the detection of ANA by ELISA (dotted blue line)

### 3.2. ANA detection in culture supernatants using commercial ANA ELISA kits

The purpose of this study was to detect ANA in culture supernatants derived from single-B cell cultures. First, we screened for the presence of ANA in culture supernatants derived from wells containing 100 cells/well and single cell sorted (SC) cells using commercial ANA ELISA kits from OriGene (ANA screen ELISA, cat# EA100913) and Eagle Biosciences (ANAscreen ELISA, cat# ANS31-K01). Neither ANA ELISA kits could detect ANA in B cell-derived culture supernatants (**Supplemental Fig. 2**).

G11 is a monoclonal IgG1 anti-double stranded (ds)DNA antibody and B1 is a IgG1 control antibody that does not bind nuclear antigens, assessed by absence of HEp2 cell staining [18]. We also tested and compared the sensitivity and detection level for B1 and G11 antibodies utilizing the same kits (**Supplemental Fig. 3**). The ANA ELISA kit from OriGene could not detect the G11 antibody even at a 400 μg/ml concentration, while the ANA ELISA assay kit from Eagle Biosciences detected G11 antibody at 10 μg/ml. These ANA ELISA kits are developed and optimized for human serum samples and not for purified antibodies or culture supernatants.

### 3.3. Development of an ANA ELISA

We developed an assay to detect ANA at low concentration by ELISA in B cell-derived culture supernatant. As demonstrated step-by-step diagram of the ANA ELISA method (**Fig. 2**). Step **A** involves coating of streptavidin on an ELISA plate overnight; step **B** involves washing the plate and adding biotinylated-nuclear extract (Bio-NE) to the ELISA plate; step **C** involves washing the plate and adding primary antibody (culture supernatant) to the ELISA plate; step **D** involves washing the plate and adding secondary antibodies (AP-conjugated); step **E** involves washing the plate and adding substrate for color development, and the final step is to monitor absorbance by ELISA reader.

**Figure 2.**
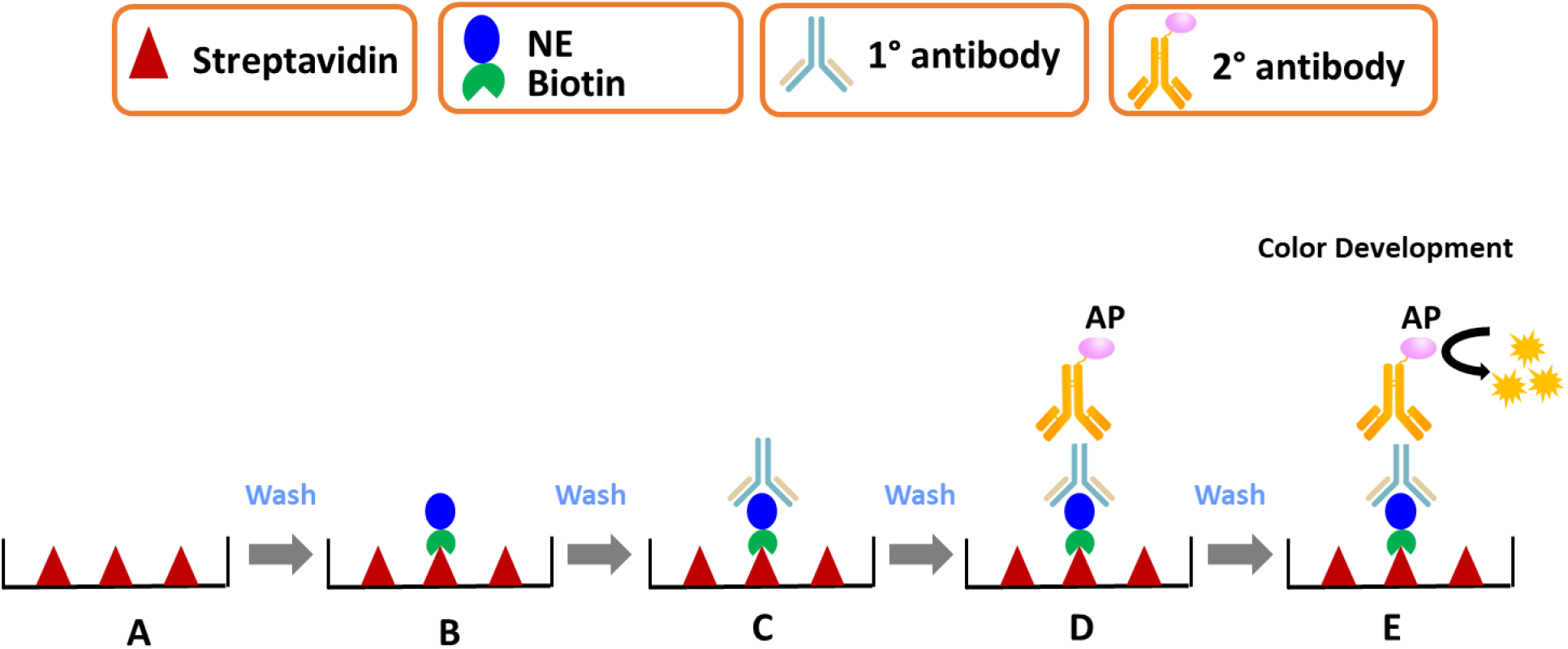
Schematic representation of the ANA ELISA method. **A**. Coating of streptavidin on ELISA plate. **B**. Addition of biotinylated-nuclear extract (Bio-NE) to the ELISA plate. **C**. Addition of primary antibodies to the ELISA plate. **D**. Addition of secondary antibodies (AP conjugated) to the ELISA plate. **E**. Addition of substrate for color development monitored by spectrophotometer.

### 3.4. Titration of ANA- and ANA+ culture supernatants (ANA-IgM ELISA)

To evaluate lowest limit of ANA-IgM, culture supernatant derived from the ANA- and ANA+ naïve B cells were titrated (1000-7.8 ng/ml). We found that the ANA-IgM ELISA can significantly detect 15.6 ng/ml in culture supernatant (**Fig. 3**). Detection of ANA+ calculation was defined based on 1.5-fold change compared to control (ANA-culture supernatant).

**Figure 3.**
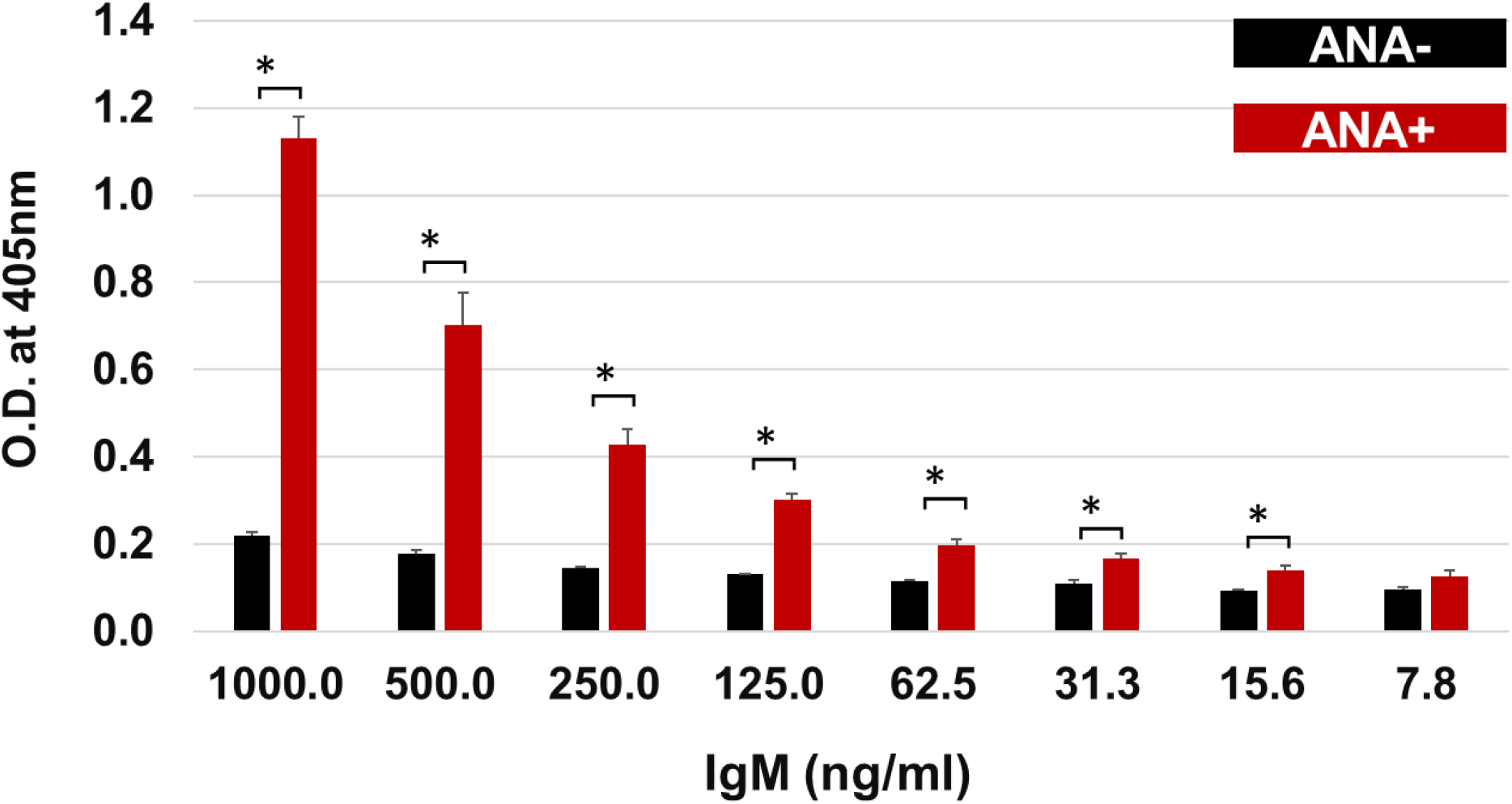
B cells culture supernatant titration (ANA-IgM). ANA ELISA assay was performed by adding serially diluted culture supernatants derived from cultures of 100 ANA- or ANA+ B cells (1000-7.8 ng/ml). Secondary antibody IgM-AP (1:500 dilution) was added, and remaining steps of ANA ELISA was performed as described. Detection of ANA+ calculation was determined based on 1.5-fold change compared to control (ANA-culture supernatant).

### 3.5. ANA detection in cultures of single sorted B cells (ANA- and ANA+ B cells)

Total Ig was estimated in samples **(Fig. 1B)** before performing ANA-Ig ELISA. The supernatants obtained from cultures of ANA-naïve B cells, which produced more than 15 ng/mL showed 2.7% (95% CI, <0.01% to 12.6%) ANA positivity (specificity of the assay = 97.3%) (**Fig. 4A**) while supernatants from ANA+ naïve B cells showed 97.8% (95% CI, 86.5% -99.7%) positivity (sensitivity of the assay) (**Fig. 4B**). This ANA ELISA assay was exclusively optimized and developed for B cell derived-culture supernatants.

**Figure 4.**
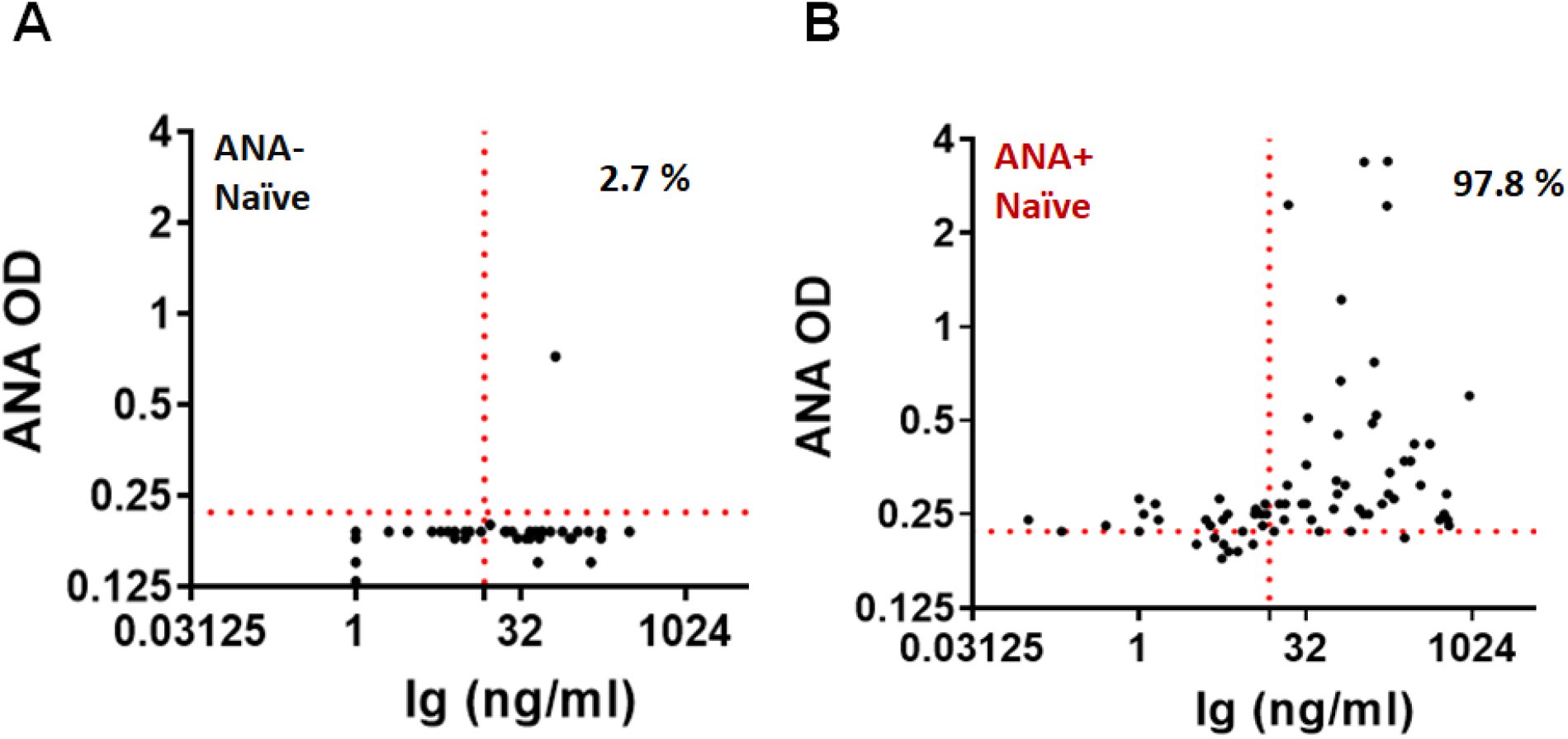
ANA detection in cultures of single sorted B cells (ANA- and ANA+ B cells). Single B cells were sorted into the 96 well culture plate. Ig production was quantified in d26 culture supernatants of ANA-naïve B cells and ANA+ naive B cells. Ig concentration >15 ng/ml considered as positive for Ig production. **A**. Detection of ANA in ANA-naïve B cell culture supernatant was performed. **B**. Detection of ANA in ANA+ naïve B cell culture supernatant was performed.

## 4. Conclusion

We have optimized a novel sensitive ELISA for the detection of ANA from single sorted naïve B cells expanded in culture for 26 days. This assay can detect ANA from as little as 15.0 ng/ml of IgM. This method will permit a comparison of the specificities of ANA produced by autoreactive B cells in healthy individuals and individuals with autoimmune disease, and should enable a study of repertoire selection in individuals with SLE and other autoimmune diseases.

## Abbreviations

ANA: anti-nuclear antibody
CS: culture supernatants
NE: nuclear extract
Bio-NE: Biotinylated-nuclear extract

