## Supplemental Figures for "Anti-nuclear antibody (ANA) detection in culture supernatant of cloned B cells"

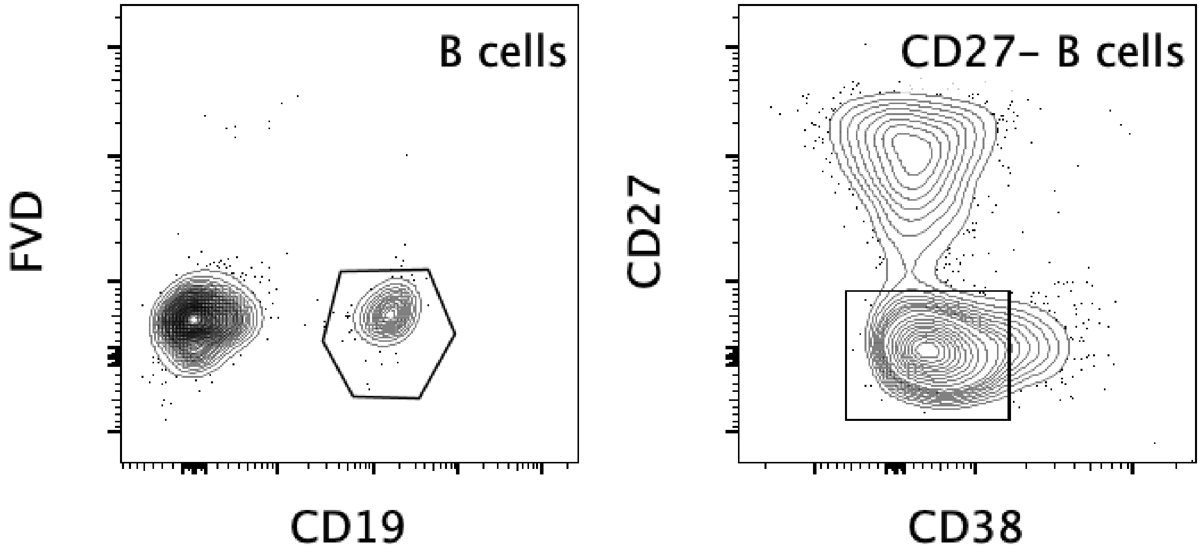


### Supplemental Figure 1

**Naïve B cell gating strategy.**

After identification of lymphocytes by FSC and SSC and single cells (not shown), live B cells were identified. Left panel shows live CD19+ B cells (CD19+, FVD-) and right panel shows naïve B cell gate (CD27-, CD38 intermediate B cells).

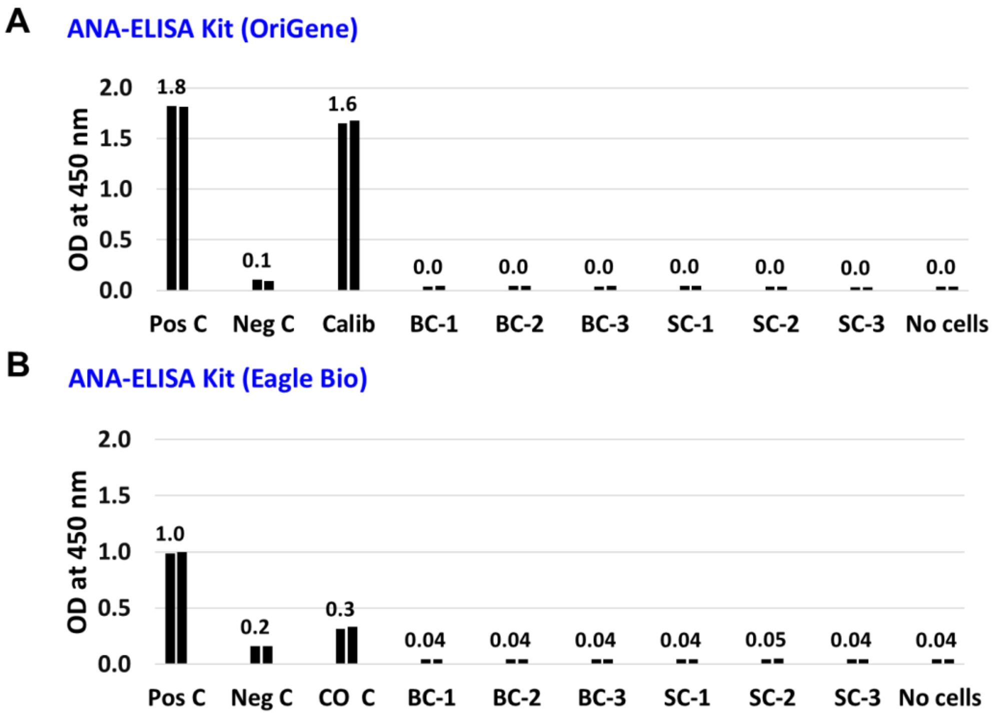


### Supplemental Figure 2

**Detection ANA in culture supernatants-derived from single cells (SC) and bulk cells (BC) using commercial ANA ELISA kits**.

**A.** ANA ELISA kit (OriGene) was used for detection of ANA in culture supernatants-derived from bulk cells (BC, 100 cells/well) and single cells (SC). ANA ELISA kit (OriGene) supplied with positive control (Pos C), negative control (Neg C), and calibrator control. **B.** ANA ELISA kit (Eagle Biosciences) was used for detection of ANA in culture supernatants-derived from bulk cells and single cells. ANA ELISA kit (Eagle Biosciences) supplied with positive control (Pos C), negative control (Neg C), and cut off control (CO C). Samples were run in duplicates as demonstrated by bars. BC: Culture supernatant from the bulk sorted cells (100 cells) and SC: Culture supernatant from the single sorted cells. The concentration of Ig within bulk cell-derived culture supernatant (~ 400 ng/ml) and single cell-derived culture supernatant (~ 150 ng/ml).

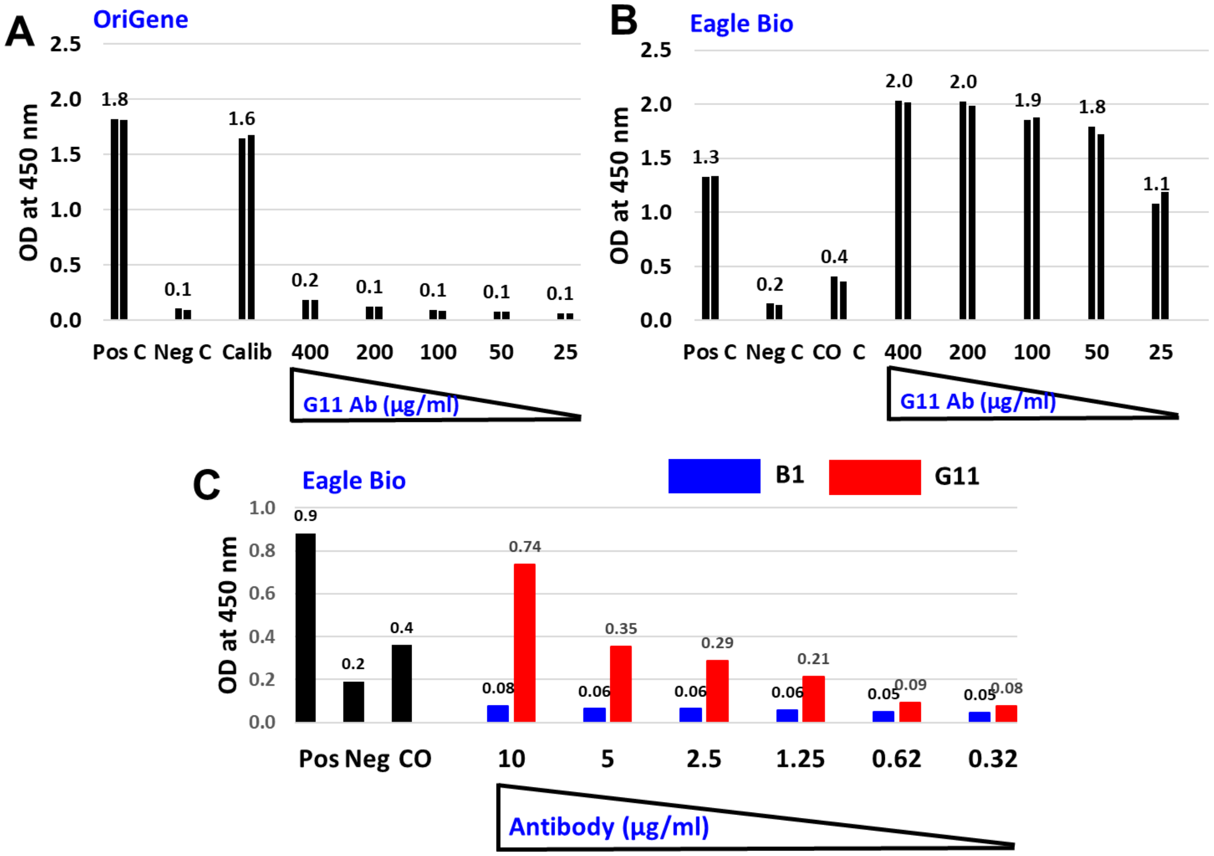


### Supplemental Figure 3

**Detection of B1 and G11 antibody was tested using commercial ANA ELISA kits.**

**A.** Titration of G11 antibody was performed by serial dilutions (400 – 25 µg/ml) using ANA ELISA kit (OriGene).

**B.** Titration of G11 antibody was performed by serial dilutions (400 – 25 µg/ml) using ANA ELISA kit (Eagle Biosciences).

**C.** Titration of both B1 and G11 antibody was performed by serial dilutions (10 – 0.32 µg/ml) using ANA ELISA kit (Eagle Biosciences). ANA ELISA kit (OriGene) supplied with positive control (Pos C), negative control (Neg C), and calibrator control. ANA ELISA kit (Eagle Biosciences) supplied with positive control (Pos C), negative control (Neg C), and cut off control (CO C). All samples were run in duplicates.
